# *ERBB2* and *KRAS* Alterations Mediate Response to EGFR Inhibitors in early stage Gallbladder Cancer

**DOI:** 10.1101/290486

**Authors:** Prajish Iyer, Shailesh V Shrikhande, Malika Ranjan, Asim Joshi, Ratnam Prasad, Nilesh Gardi, Rahul Thorat, Sameer Salunkhe, Bhasker Dharavath, Bikram Sahoo, Pratik Chandrani, Hitesh Kore, Bhabani Mohanty, Vikram Chaudhari, Anuradha Choughule, Dhananjay Kawle, Pradip Chaudhari, Arvind Ingle, Shripad Banavali, Mukta R Ramadwar, Kumar Prabhash, Savio George Barreto, Shilpee Dutt, Amit Dutt

## Abstract

The uncommonness of gallbladder cancer has contributed to the generally poor understanding of the disease, with scant reports restricted to advance-stage tumors. Here, using an integrated analysis of whole exome and phospho-proteome, we show recurrent activating *ERBB2* and *KRAS* somatic mutations are present in 6 and 3 of 44 early-stage rare gallbladder tumors, respectively. *In vitro* and *in vivo* cell-based and biochemical assays reveal an essential role of ErbB pathway activation for the survival of gallbladder cells. Interestingly, the genetic and pharmacological dependencies of gallbladder cells are dependent on the *KRAS* mutant allele status, reminiscent of the clinical algorithm commonly practiced to opt for anti-EGFR treatment in colorectal cancer. In overall, we present the first evidence that the presence of *KRAS* (G12V), but not *KRAS* (G13D) mutation, may preclude gallbladder cancer patients to respond to anti-EGFR treatment, leading to an early adoption of an approved treatment regimen for gallbladder cancer patients.

## Introduction

Genomically matched therapies targeting activated tyrosine kinases have shown promise across multiple cancer types (Krause & Van Etten, 2005). The success of tyrosine kinase inhibitors (TKIs) such as imatinib, a BCR-ABL fusion protein inhibitor (Druker et al, 2001); vemurafenib, a RAF inhibitor (Flaherty et al, 2010); lapatinib, an inhibitor of ERBB2 (Rusnak et al, 2001); erlotinib and crizotinib, inhibitors of EGFR and ALK, respectively (Christensen et al, 2007; Paez et al, 2004); and, others have provided a powerful validation for precision cancer medicine. Although these treatments offer great promise, selective genomic profiling of tumors tend to impede broader implementation of genome-based cancer care (McGranahan & Swanton, 2015). For example, an inadequacy to account for multiple relevant genetic alterations likely resulted in comparable outcomes in a recently performed randomized trial where multiple cancer type patients were profiled for selected driver alterations and randomized to receive genomically-matched versus conventional therapy (Le Tourneau et al, 2014). Such important clinical studies underscore the need for convergence of information for multiple genetic alterations to ensure success of future clinical trial designs, with specific emphasis for consideration of co-occurring alterations that could potentially render tumors unlikely to benefit from genomically-matched treatments. Some prototypical examples include *KRAS*, *NRAS*, and *BRAF* mutations in colorectal cancers or secondary *EGFR* mutations in lung cancer against anti-EGFR targeted therapies (De Roock et al, 2010b).

The EGFR family of receptor tyrosine kinases (RTK) consists of *EGFR*, *HER2*, *HER3* and *HER4* (human EGFR-related-2, -3, and -4). A ligand-bound EGFR family member forms a homo- or hetero-dimer to activate the PI3K-AKT-mTOR or RAS-RAF-MAPK downstream signaling pathway to evade apoptosis and enhance cell proliferation (Garner et al, 2013). Interestingly, of all EGFR family members, HER2 lacks a ligand binding domain and forms preferred partner for other members to heterodimerize even in the absence of ligand (Spivak-Kroizman et al, 1992). Deregulation of EGFR family RTK-signaling network endows tumor cells with attributes to sustain their malignant behavior and survival, as is frequently observed in breast cancer, lung cancer, pancreatic cancer, head and neck cancer and colorectal cancer (Tebbutt et al, 2013). Interfering with the EGFR pathway thus forms the basis for the development of targeted anticancer therapies such as RTK-targeted antibodies (Cetuximab and Herceptin) and small-molecule inhibitors of RTK kinase (Erlotinib, Lapatinib, Afatinib, etc.) that have shown dramatic clinical response (Tebbutt et al, 2013). In such responses, however, co-occurrence of a *KRAS* mutation – a downstream component of the pathway-preclude patients from anti-EGFR treatment in colorectal cancer, wherein *KRAS* codon 12, but not codon 13 mutations are associated with poor outcomes (Yoon et al, 2014), underscoring their prognostic impact.

Gallbladder cancer, the most common malignancy of biliary tract, is a rare form of cancer in the world where chemotherapy and other palliative treatments have little effect on overall survival of patients (Rakic et al, 2014). The poor understanding of gallbladder cancer due to its uncommonness in the western world but high prevalence in Chile and the Indian subcontinent lends itself to the need for further research (Barreto et al, 2014). While the 5-year survival rate of an early stage T1 gallbladder carcinoma is nearly 100%, it signficantly decreases as the disease progresses, with less than 15% for T3/T4 advanced stage tumors(Zhu et al, 2010). A hope for longer term survival has specifically been promising for an early stage T2 carcinomas with an intermediate 5-year suvival (Miller & Jarnagin, 2008). Literature suggests *HER2* overexpression in 12–15 % of advanced stage gallbladder cancers with a favorable response to HER2 directed therapy (Javle et al, 2015). Moreover, three recent studies analyzed whole exome sequence of advanced stage gallbladder tumors with consistent findings (Barreto et al, 2014; Jiao et al, 2013; Nakamura et al, 2015). In order to understand the landscape of somatic alterations among a clinically distinct early staged pT1/pT2 gallbladder cancer patients, we performed whole exome sequencing of 17 early staged tumor-normal paired gallbladder samples, 5 gallbladder cancer cell lines followed by validation in 27 additional tumor samples. Here, we report novel somatic mutations of *ERBB2* in gallbladder cancer, and its therapeutic implication in presence and absence of *KRAS* (G12V) and (G13D) mutations.

## Results

### Integrated genomics and proteomics approach identify aberrant alterations in members of the *EGFR* family in gallbladder cancer

We performed whole-exome sequencing on paired tumor and germline DNA samples from 17 patients with gallbladder cancer and 5 gallbladder cancer cell lines (Supplementary Table S1 and S2). We achieved >100-fold mean sequence coverage of targeted exonic regions. The average non-synonymous mutation rate was found to be 7.7 mutations per megabase (Supplementary Table S3), which is significantly higher than as reported for other populations (Li et al, 2014). The nucleotide mutation pattern was observed to be enriched for C>T transition followed by A>G transition (Supplementary Figure S1), consistent with previous reports (Li et al, 2014). A total of 5060 somatic variants found across 17 tumors consisted of 3239 missense, 1449 silent, 131 nonsense, 135 indels and 106 splice site mutations. Somatic mutations in genes previously reported to be altered in gallbladder cancer, including recurrent mutations in *TP53* (35.2%)*, ERBB2, SF3B1, ATM* and *AKAP11* at 17.6% each were found to be mutated at comparable frequencies (Li et al, 2014) (Figure 1A and Supplementary Table S3). For validation of a few *TP53, ERBB2, ERBB3, SMAD4* and *CTNNB1* mutations, sanger-based sequencing were carried out in a subset of patients (Supplementary Figure S2). Among set novel alterations, we observed significant somatic mutations in chromatin modifier genes such as *SF3B1*, *ATRX*, *CREBBP* and *EZH2* that are known to play a significant role in other cancer types (Yoshida & Ogawa, 2014). In addition, we also found two tumor samples that harbored known activating kinase domain mutations in *ERBB2*, (V777L) and (I767M) (Bose et al, 2013); while two samples harbored *EGFR* (I1005V) and *ERBB3* (R112H) mutation (Supplementary Table S4). We identified 5 more samples with *ERBB2* mutations harboring (V777L) mutations in an additional set of 27 gallbladder cancer samples (Figure 1B). Interestingly, copy number analysis using cghMCR software identified *EGFR* amplification with a highest Segment Gain Or Loss (SGOL) score of 18 (Figure 1A), as reported earlier (Javle et al, 2015). In overall, we observed genomic amplification in *EGFR, CDK4, MDM4, CCND1, CCNE1, MYC, STK11* and *BRD3,* and deletion in *FHIT*, *SMAD4*, *TRIM33* and *APC*.

**Figure 1:**
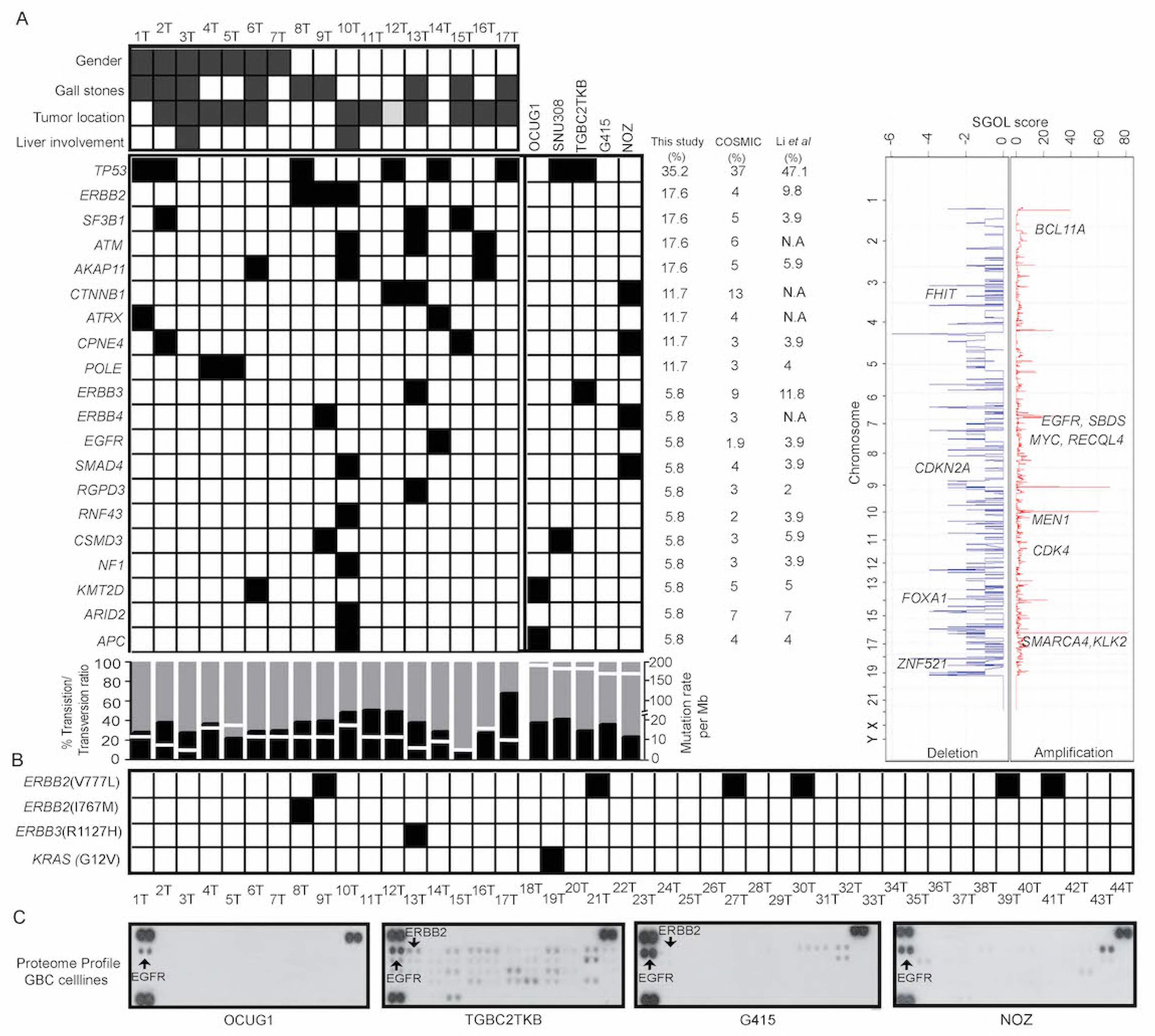
Integrated genomic and proteomic analysis of gallbladder cancer. **A)** The heat map represents somatic mutation landscape in gallbladder cancer patients (n=17) and primary tumor derived cancer cell lines (n=5) using whole exome sequencing. Clinicopathological features such as gender, gallstones, tumor location and liver involvement are shown for each patient. The grey solid boxes denote females, presence of gallstone, tumor location (neck) and positive for liver involvement. The white box denotes males, absence of gallstones, tumor location (body) and negative for liver involvement. The genes are arranged in the decreasing order of their frequency. Black solid box indicates the presence of mutation in the heatmap. Mutation frequencies of the genes mentioned are shown in this study, COSMIC-GBC and Li *et. al* study. The transition to transversion ratio is shown in percentage for each patient indicated by different shades (Black denotes transversion and grey denotes transition). Somatic mutation rate/30Mb is derived from whole exome sequencing data is indicated by white line. Overall copy number changes derived from whole exome sequencing data. The horizontal-axis is represented by a score of segment gain or segment loss (SGOL score) while the vertical-axis represents the chromosomal positions. Copy number gain is indicated by red with positive SGOL score while copy number loss is indicated by blue with a negative SGOL score. Representative cancer-associated genes are annotated in their respective amplified/deleted regions. **B)** Schematic representation of *ERBB* family mutation validation by Sanger sequencing in an additional set of 27 samples. Solid box indicate presence for mutation in the respective samples, white boxes indicates no event. **C)** RTK array analysis of gallbladder cancer cells (OCUG1, TGBC2TKB, G415, and NOZ) for 10 minutes exposure of blot is shown. Each RTK is spotted in duplicate and the pair of dots in each corner of the membrane corresponds to positive and negative control. Tyrosine phosphorylation of EGFR (ERBB1) and ERBB2 were observed consistently, indicated by arrow.

Next, to correlate differential activation of signaling molecules with their genomic alterations, we performed phospho-proteomic profile of four gallbladder cell lines for 49 receptor tyrosine kinases using a phospho-RTK array. Consistent with whole exome findings, we observed varying levels of EGFR and ERBB2 constitutive phosphorylation in all gallbladder cancer cell lines based on their phospho-proteome (Figure 1C) and follow up validation by western blot analysis (Supplementary Figure S3A). Interestingly, the whole exome data analysis and Sanger sequencing based validation also revealed that 1 of 44 gall bladder patients and NOZ cells harbor *KRAS* (G12V) mutation; G415 cells harbor *KRAS* (G13D) mutant allele; while OCUG1 and SNU308 cells were wild type for *KRAS* (Figure 1B; Supplementary Figure S2). These four cell lines thus represent diverse gallbladder cancer sub-classes based on their *KRAS* mutant allele status (Li et al, 2014). Of note, *KRAS* mutations are known to predict plural clinical outcome in response to EGFR inhibitors in colorectal and lung cancer along with other mutations (Supplementary Table S4) (Choughule et al, 2014).

### *ERBB2* and *EGFR* are essential for gallbladder cancer cells not harboring *KRAS G12V* mutant allele

To determine the significance of *EGFR* and *ERBB2* constitutive phosphorylation and *KRAS* mutant alleles in gallbladder cancer cells, we set out to establish whether expression of *ERBB2* is required for gallbladder tumor cell survival. We tested a series of five shRNA constructs in three gallbladder tumor cell lines expressing *ERBB2* with wild type *KRAS* in OCUG1 cells, along with G415 and NOZ cells harboring the *KRAS* (G13D) and *KRAS* (G12V) mutant alleles, respectively. We identified three shRNA constructs that efficiently knocked down expression of *ERBB2* and inhibited the constitutive phosphorylation of MAPK in OCUG1 and G415 cells but not in NOZ cells (Figure 2A), consistent with drug sensitive outcome described in colorectal cancer wherein cells harboring wild type *KRAS* or mutant *KRAS* (G13D) allele are sensitive to *EGFR* inhibitor but not those harboring mutant *KRAS* (G12V) mutant allele (Osumi et al, 2015). This suggests that *KRAS* (G13D) but not *KRAS* (G12V) still requires upstream *EGFR* signaling in gallbladder cancer cells, similar to as established in colorectal cancer(Kumar et al, 2014). Next, we used these cells to demonstrate that knockdown of *ERBB2* inhibited anchorage-independent growth, cell survival, cell invasion and migration efficiently in OCUG1 and G415 cells but not in NOZ cells (Figure 2B-E). Furthermore, as unlike other *EGFR* family members, *ERBB2* does not require ligand binding for dimerization but can be activated by heterodimerization (Linggi & Carpenter, 2006), we asked if *EGFR* mediates the activation of downstream signalling pathways. We performed co-immunoprecipitation of *EGFR* and *ERBB2* to establish that *ERBB2* interacts with *EGFR* in gallbladder cells (Supplementary Figure S3B), possibly similar to *ERBB3* as shown earlier in gallbladder cells (Li et al, 2014). Moreover to test if *ERBB2* requires *EGFR* also for sustained signaling and transforming potential, we knocked down the expression of *EGFR* in OCUG1 and G415 cells. The knockdown of *EGFR* inhibited anchorage-independent growth, cell survival, cell invasion and migration in OCUG1 but not in G415 cells, similar to *ERBB2* knockdown (Supplementary Figure S4). Taken together, this suggests that *ERBB2* requires *EGFR* or other members of the family possibly to dimerize for activation, such that down-regulation of *EGFR* and potentially other members suppress the functionality of *ERBB2,* as has been previously reported in breast cancer (Zhou & Agazie, 2012).

**Figure 2:**
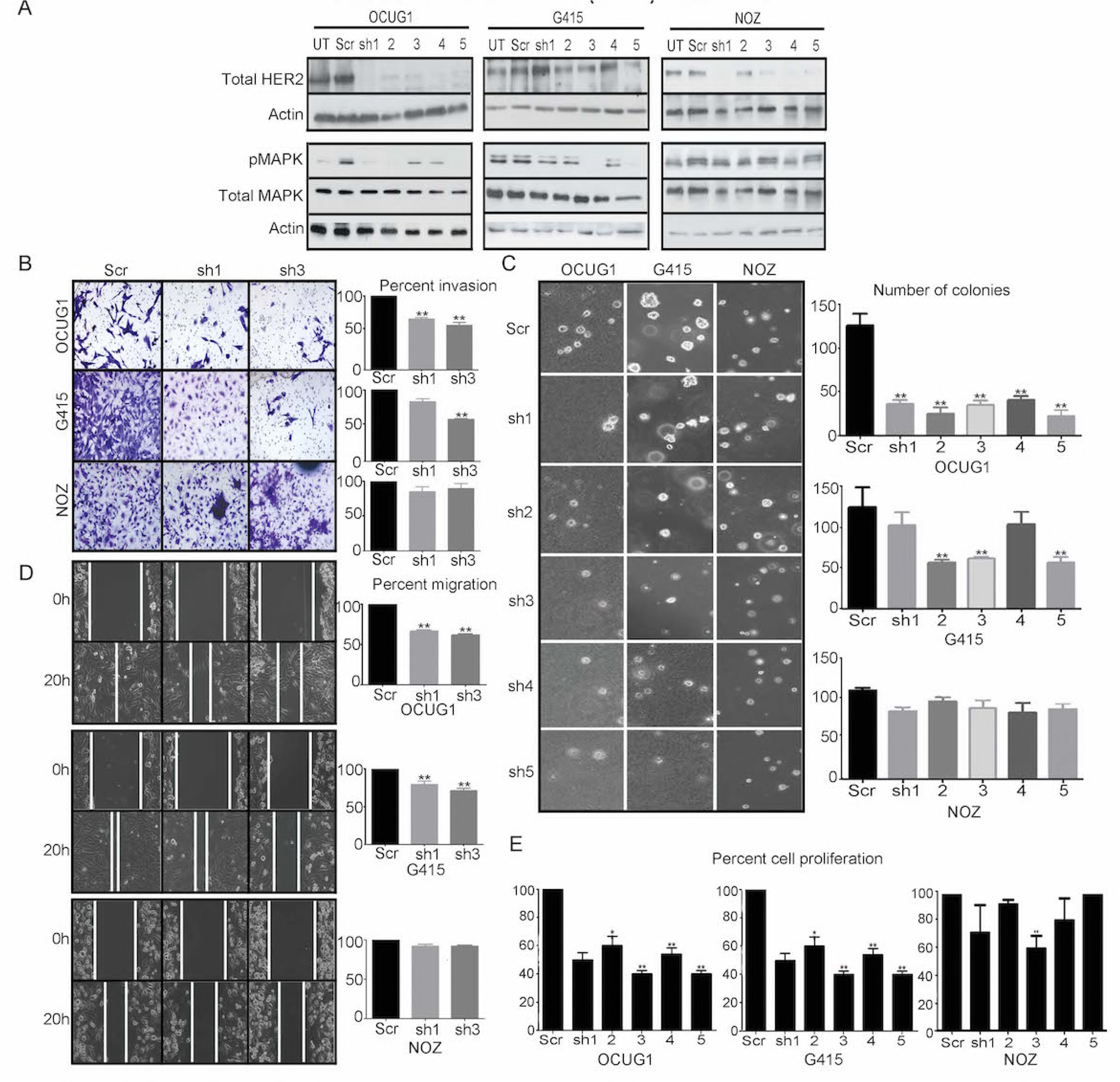
Knockdown of *ERBB2* expression with shRNA inhibits survival of gallbladder cancer cells that do not harbor *KRAS* (G12V) mutant allele. **A)** Western blot analysis with 5 shRNA constructs used to knock down ERBB2 expression were packaged into lentivirus and used to infect OCUG1, G415, and NOZ cells. Anti-ERBB2 immunoblot shows that hairpins 3 and 5 efficiently and consistently knock down endogenous *ERBB2* expression across all cells (**A upper panel**) with concomitant decrease in downstream signaling as assessed by anti-phospho-MAPK immunoblot in OCUG1 and G415 cells but not in NOZ cells that harbor a constitutivelyactive KRAS (G12V) mutation (**A lower panel**). Actin is included as a loading control. Scr, scrambled hairpin and untransfected cells (UT) used as a negative control. Knockdown of *ERBB2* expression with shRNA inhibits; invasion characteristics as assessed by matrigel assay (**B**); anchorage-independent growth as shown by soft agar assay(**C**) and, migration as assessed by scratch assay (**D**) of OCUG1 (with wild type *KRAS*) and G415 (with *KRAS* (G13D)) cells but not NOZ gallbladder cancer cell lines that harbor an activating *KRAS* (G12V) mutation. The graph on the right panel represents percent inhibition normalized to scrambled (Scr) control cells. Similarly, knockdown of *ERBB2* expression with shRNA inhibits percent growth as determined by MTT assay with bar graph plotted with readings obtained on day 4 relative to day 1 for OCUG1, G415, and NOZ cells **(E)** for each shRNA construct and normalized to scrambled control cells. Representative plates from three independent experiments are presented. Colonies were photographed and quantitated after 2 weeks for soft agar assay (Magnification: ×10); 1 day for invasion; and 20 h for migration assay. *P<0.05.

### Gallbladder cancer cells not harboring *KRAS* (G12V) mutant allele are sensitive to irreversible *EGFR* inhibitors *in vitro* and *in vivo*

Next, we investigated whether inhibition of kinase activity of EGFR family receptor tyrosine kinases would be effective against gallbladder cancer cell lines. Treatment of the OCUG1 and G415 cells with BIBW-2992 (Li et al, 2008), but not reversible EGFR inhibitor gefitinib (data not shown), similarly abolished phosphorylation of MAPK in OCUG1 cells, which was constitutively phosphorylated in the untreated gallbladder cell lines compared to the NOZ cells and resulted in a marked decrease in migration, invasion, colony formation in soft agar and cell survival in liquid culture, with IC50s of 0.8 uM in OCUG1 and 2.0 uM in G415 cells, whereas no effect was observed on NOZ cells harboring *KRAS* (G12V) mutant allele (Figure 3A-E). Furthermore, when injected subcutaneously into NOD/SCID mice, 13 of 13 mice injected with G415 cells formed tumors ∼13 days post injection; 10 of 10 mice injected with NOZ cells ∼6 days of post injection; while none of 10 mice injected with OCUG1 cells formed tumors uptill 2 months post injection of cells (Supplementary Table S8). When the tumors reached ∼100-150mm^3^, tumors were treated orally with 15mg/Kg irreversibile EGFR inhibitor Afatinib- or vehicle for a period of 15 days. Consistent with *in vitro* data, tumors treated with Afatinib slowed or reversed their growth compared to vehicle in G415 xenografts (n=7) but not in NOZ (n=6) xenografts. The overall effect on tumor burden in vehicle-treated versus Afatinib-treated mice were 5.7-folds lower in G415 xenografts, while no significant differences were observed in in NOZ xenografts(Figure 4A and B). This reduction in tumor size in G415 xenografts was paralleled by reduction in the amounts of phospho-ERK1/2 by immuno-histochemical analyses (Figure 4C-D, lower panel) of explanted tumors, further validating our *in vitro* findings (Figure 3A).

**Figure 3:**
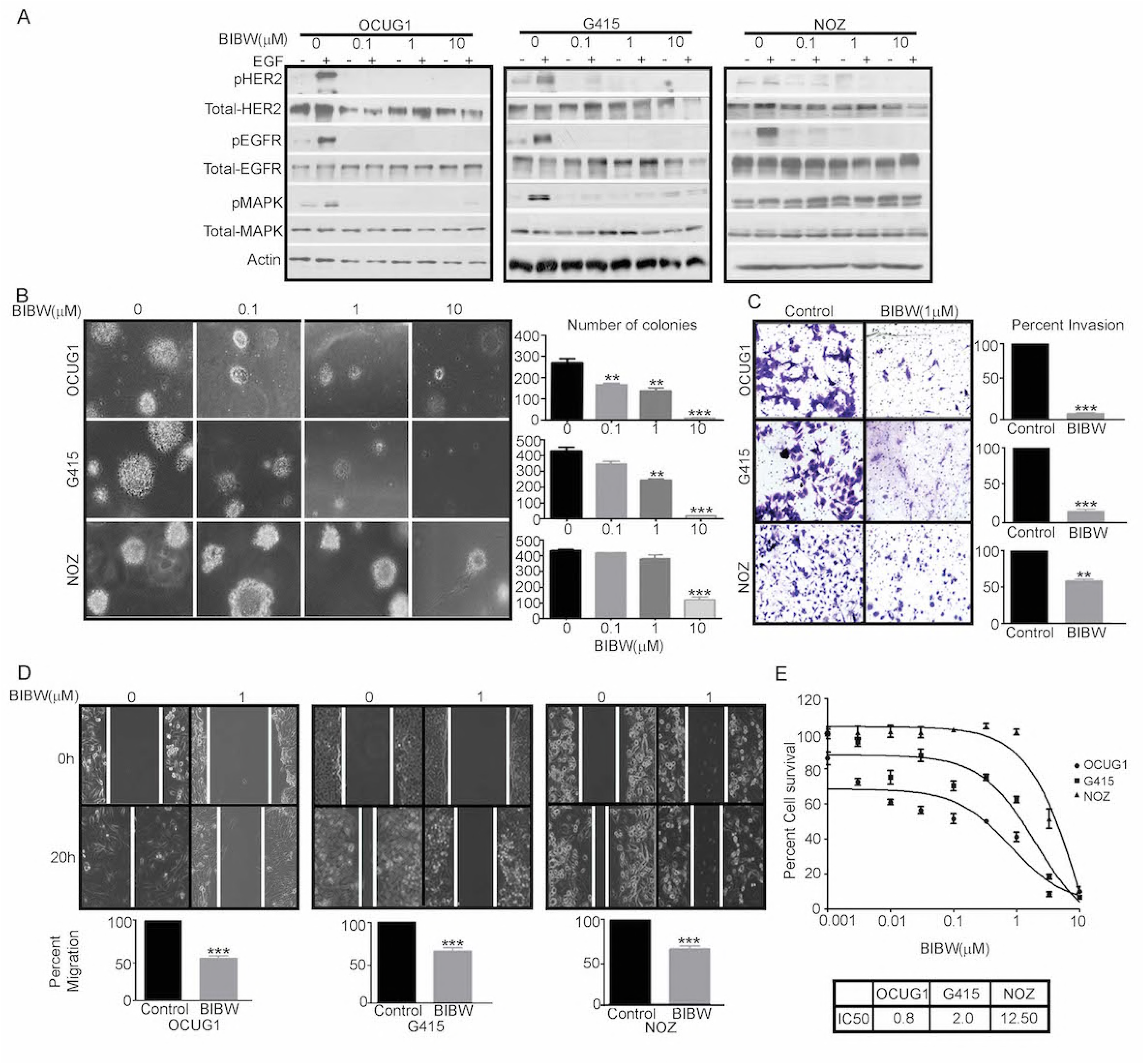
ERBB2 tyrosine kinase activity is essential for gallbladder cancer cells that do not harbor *KRAS* (G12V) mutant allele. **A)** Treatment of OCUG1, G415 and NOZ gallbladder cancer cells for 10-12 hrs with 0-10 µM covalent EGFR inhibitor BIBW-2992 inhibits both basal and ligand-induced (5-min stimulation with 20 ng/ml EGF) EGFR and ERBB2 phosphorylation, as evident from immunblotting with anti-phospho antibodies specifically recognising EGFR (pY1068) and ERBB2 (pY1248). However, EGFR inhibitor BIBW-2992 inhibits MAPK activation as determined by pMAPK p42/p44 (Thr202/Thr204) antibody, a downstream effector component of EGFR- and ERBB2-dependent signalling pathways in OCUG1 (with wild type *KRAS*) and G415 (with *KRAS* (G13D)) cells but not in NOZ gallbladder cancer cell lines that harbor an activating *KRAS* (G12V) mutation. Actin was used as a loading control. Treatment with the indicated concentrations of EGFR inhibitor BIBW-2992 inhibited soft agar colony formation (**B**); invasion (**C**); and, migration (**D**) by the OCUG1, G415 but not NOZ gallbladder cancer cell lines with hyper phosphorylated ERBB2. *P<0.05 vs control. Representative plates from three independent experiments are presented. Colonies were photographed and quantitated after 2 weeks for soft agar assay (Magnification: ×10); 1 day for invasion; and 20 h for migration assay. Quantification of effects of BIBW-2992 for assays is indicated in the form of bar graph. **D)** Gallbladder cancer cell line MTT survival assays performed after 72h treatment with BIBW-2992 in 6 replicates is shown. IC_50_s are indicated in form of table as determined by nonlinear regression with Prism GraphPad software.

**Figure 4:**
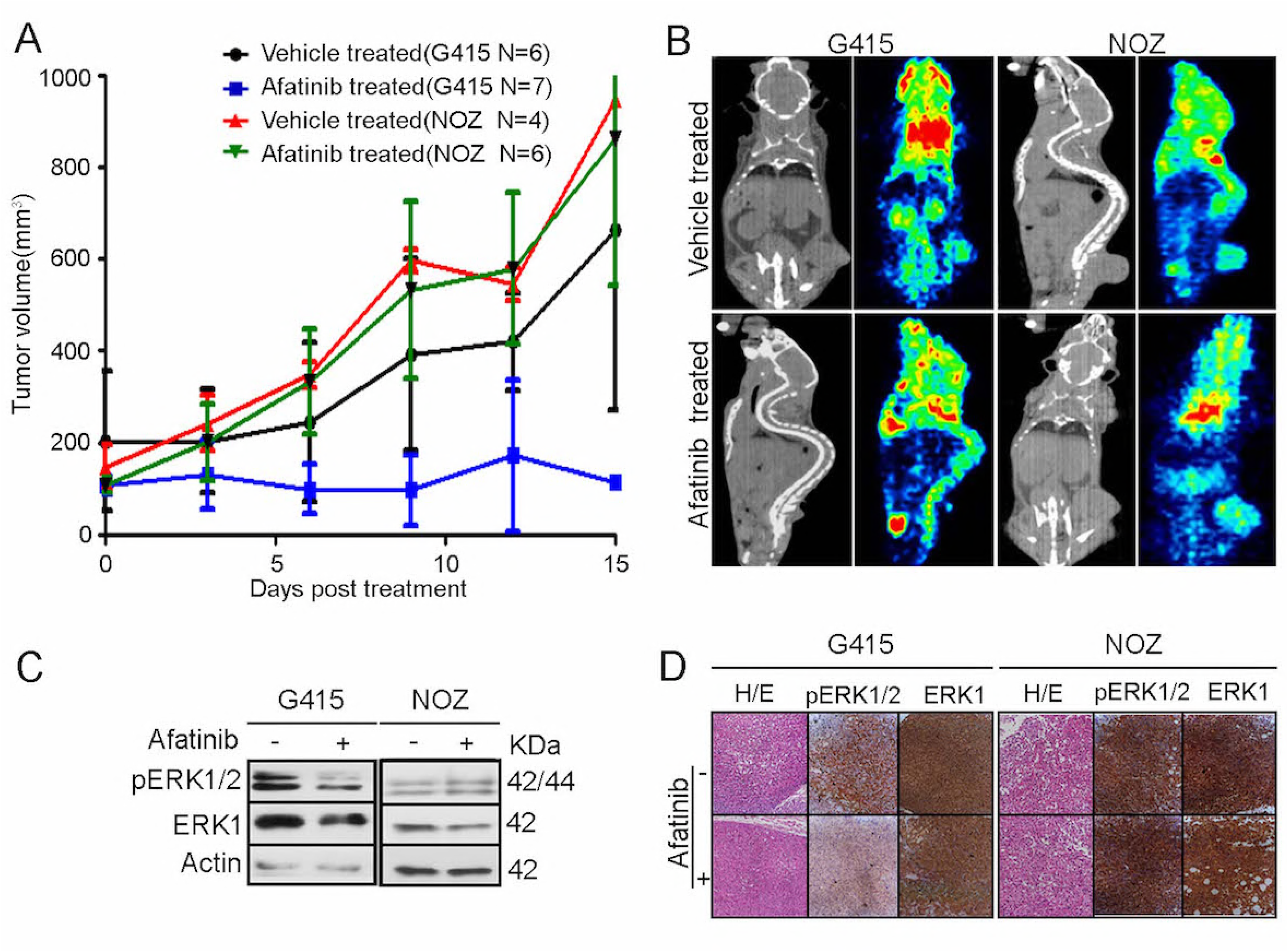
*In vivo* sensitivity of gallbladder cancer cell lines to EGFR inhibitor. **A)** G415 and NOZ xenografts developed in NOD-SCID mice were subjected to afatinib (15mg/kg) or vehicle treatment for a period of 15 days. The plot shows the tumor volume (mm^3^) during the course of drug treatment indicating reduction of tumor volume in afatinib treated G415 xenografts. **B)** CT scan and PET imaging by F^18^-FDG uptake is shown for vehicle and afatinib treated xenografts. The gradient color code is shown for uptake of F^18^-FDG with red indicating maximum uptake **C)** Immunoblot analysis of phosphorylation of MAPK (pERK1/2, ERK1) is shown for vehicle(-) and afatinib(+) treated xenografts. Actin is used as the loading control. D) Immuno-histochemical staining of pERK1/2, ERK1 is shown for vehicle(-) and afatinib(+) treated xenografts.

### Clinical correlation of *TP53* and *EGFR* family mutations in early stage gallbladder cancer patients

In overall, the patient cohort represents a good subset of fairly early stage disease who received experienced and good quality radical surgery in a tertiary referral center. The Kaplan-Meier survival analysis with respect to *TP53* mutation status revealed an overall survival of 40 months (n=6; 95% CI: 34.1-70.7) compared to 52 months (n=11; HR: 0.8, 95% CI: 1.1-4.4 P= 0.799) among patients with wild type *TP53* (Supplementary figure 5A). Thus, the overall survival of patients with mutations decreased as compared to patients with wild type *TP53*. This observation is consistent with previous reports wherein *TP53* mutations are associated are known to be associated with worse prognosis in various cancer types (Kandioler-Eckersberger et al, 2000). Interestingly, overall survival with respect to *EGFR* family mutation status was 65 months (n=9; 95% CI: 49.5-.82.1) compared to 54 months (n=35; 95% CI: 42.1-65.2) among patients with wild type *EGFR* family genes (Supplementary figure 5B; Supplementary table S5).

## Discussion

This study represents the first genomic landscape of an early-stage gallbladder cancer that reveals somatic mutations in *TP53, ERBB2, ATM, AKAP11, SMAD4* and *CTNNB1* similar to as reported in advance-stage gallbladder tumors. Our mutation pattern analysis revealed an enrichment for C>T transition followed by A>G transition, a signature which suggests an underlying chronic inflammation leading to GC to AT polyclonal transition (Yanagisawa et al, 2010), as reported earlier (Iyer et al, 2016). We also observed significant somatic mutations in chromatin modifier genes such as *SF3B1*, *ATRX*, *CREBBP* and *EZH2* that have not been reported earlier in gallbladder cancer, indicating potential therapeutic options. Analyzing the potential effects of somatic alterations on survival of gallbladder cancer patients, we observed a trend among patients with wild type *TP53* to survive longer than patients with *TP53* mutations, which is known to predict failure of chemotherapy in several cancer types (Petitjean et al, 2007) and is consistent with previous reports observed in gallbladder cancer (Doval et al, 2014).

Additionally, consistent with a recent report that described alterations in *ERBB2* and *ERBB3* at a frequency of 9.8% and 11.8% respectively among Chinese gallbladder cancer (Li et al, 2014), we found recurrent activating *ERBB2* (V777L) mutation in 6 of 44 gallbladder cancer samples with an overall mutation frequency of 13%, in addition to *ERBB3* (R112H) and *EGFR* (I1005V) mutation occurring at 2%, each in our sample set. The (V777L) alteration has been shown to be sensitive to lapatinib in biliary tract cancer, breast cancer cell lines and other isogenic systems overexpressing the alteration (Bose et al, 2013; Javle et al, 2015). Functional studies performed using gallbladder cell lines establish that *ERBB2* and *EGFR* are essential for the survival of gallbladder cancer cells. Given that *ERBB2* lacks the ligand binding domain, the co-immunoprecipitation experiments suggest that ERBB2 dimerize with EGFR, and possibly with other members, to constitutively activate the pathway. Interestingly, genetic or pharmacological ablation of *ERBB2* and *EGFR* function, using EGFR small-molecule irreversible inhibitor BIBW-2992, diminishes the survival, anchorage-independent growth, migration and invasion characteristics of gallbladder cancer cell lines, suggesting members of the EGFR family as an effective therapeutic target. Furthermore, while *KRAS* mutations in gallbladder cancer have been reported to occur at a frequency from 3% to 30 % (Muller et al, 2014), some co-occuring with activating *ERBB3* mutation, we observed *KRAS* (G12V) and (G13D) mutation in 1 of 44 primary gallbladder tumors and 2 of 5 gallbladder cancer cell lines that are known to be associated with differential clinical outcome in response to anti-*EGFR* therapy in colorectal cancer (De Roock et al, 2010a; Li et al, 2015). The biological characteristics of *KRAS* mutation is known to vary by cancer types as those found in pancreatic and non-small cell lung cancers are predominantly at codon 12, while in colorectal and gallbladder mutations appears to be in codon 12 and codon 13 (Prior et al, 2012). Moreover, clinical response among patients along with *invitro* and *invivo* studies with isogenic colon cell line indicate *KRAS* (G13D) mutation as sensitive but (G12V) as resistant to anti-*EGFR* therapy suggesting codon 13 mutations are still dependent on inductive upstream *EGFR* signaling and exhibit weaker *in vitro* transforming activity than codon 12 mutations (De Roock et al, 2010a).

In summary, besides suggesting adoption of anti-EGFR therapy as a therapeutic option in early-stage gallbladder cancer based on *ERBB2* alteration, we present the first evidence that presence of *KRAS* (G12V) but not *KRAS* (G13D) mutation may preclude such patients to respond to the treatment, similar to the clinical algorithm commonly practiced based on *EGFR* alteration in colorectal cancer. As low prevalence rate of the disease, target accrual in clinical trials has been a bottleneck in gallbladder cancer, this study forms the basis to include gallbladder patients for an anti-EGFR therapy under basket clinical trials such as the NCI–Molecular Analysis for Therapy Choice (NCI-MATCH) trials that are genomically matched (Do et al, 2015).

## Materials and Methods

### Patient Information

A total of 27 fresh frozen samples (10 tumor-normal paired and 7 orphan tumors) were utilized for whole exome sequencing. An additional set of 27 FFPE samples were utilized as a validation set. Tumor-normal paired samples were collected at Tata Memorial Hospital and Advanced Centre for Treatment, Research and Education in Cancer (ACTREC), Mumbai. Sample set and (ACTREC-TMC) Internal Review Board (IRB) -- IRB Project Number # 104--approved study protocols. Formalin-fixed paraffin-embedded tissue blocks were collected from the tissue repository of Tata memorial hospital (TMH-TTR) in compliance with the guidelines. These tissues were examined for tumor content and the tumor content was in the range of 40-90%. Patient samples and characteristics are provided in the Supplementary Table S1 and S5.

### DNA extraction

Genomic DNA was extracted from fresh frozen samples by using Qiagen Blood and Cell culture DNA kit. The extracted DNA yield and quality were assessed using Nanodrop ND2000 (Thermo scientific). The extracted DNA (about 1µg) from the fresh-frozen tissue specimens were sent to Genotypic Technology Pvt Ltd, Bangalore for exome sequencing. Genomic DNA from FFPE blocks was extracted using Qiagen QiAmp DNA FFPE Tissue kit as per manufacturer instructions. The extracted DNA yield and quality were assessed using Nanodrop ND2000 (Thermo scientific). These samples were further checked for integrity by PCR amplification of GAPDH (96bp). These samples were used for extended Sanger validation of identified variants in exome sequencing.

### Exome analysis pipeline and somatic mutation calling

The variant analysis was performed as described previously (Chandrani et al, 2017; Upadhyay et al, 2016a). MutSigCV v2.0 (Lawrence et al, 2013) and IntOgen (Gonzalez-Perez et al, 2013) were used for identification of the significantly mutated gene and p value ≤0.05 was considered as the threshold for significance. The variants were excluded if they were present in exclusively in dbSNP, TMC-SNPdb or both. Also, we removed variants that were identified in all three databases – COSMIC (v68) (Forbes et al, 2008), dbSNP (v142) (Sherry et al, 2001) and TMC-SNPdb database (Upadhyay et al, 2016a). The annotated cancer-associated variants were annotated using Oncotator (v1.1.6.0) (Ramos et al, 2015) and restricted our further analysis to only coding variants. Intogen (https://www.intogen.org/search) was used to calculate the significance of frequently mutated gene in our cohort. Since our dataset was inherently not suitable for above tools due to limited number of tumor samples (n=17), we have also performed extensive functional prediction tool based analysis for non-synonymous variants using nine different tools as described earlier (Chandrani et al, 2017). Total number of identified somatic substitutions in exome sequencing was extracted from MutSigCV output and was processed to calculate the number and frequency distribution of various transitions and transversions.

### Exome sequencing capture, library construction, and sequencing

Exome capture and sequencing were performed as described previously (Upadhyay et al, 2016b). Briefly, Agilent Sure select in-solution (low-input capture-500ng) were used to capture ∼62Mb region of human genome comprising of ∼201,121 exons representing ∼20,974 gene sequences, including 5’UTR, 3’UTR, microRNAs and other non-coding RNA. Sequencing was run with 150bp paired end reads to achieve coverage of 100X and was performed according to Illumina standard protocol.

### Copy number analysis from Exome sequencing data

Control-FREEC (Boeva et al, 2012) was used for copy number analysis from BAM files of variant calling analysis. Genes with Segments-of-Gain-Or-Loss (SGOL) score ≥4 were defined as amplified genes and ≤-2 as deleted genes by cghMCR package of R (http://bioconductor.org/packages/release-/bioc/html/cghMCR.html). The validation of somatic copy number changes was performed as described previously (Upadhyay et al, 2016b).

### Cell culture and reagents

Human GBC cell lines (OCUG1, SNU308, TGBC2TKB, NOZ, and G415) obtained as a kind gift from Dr. Akhilesh Pandey (IOB, Bangalore) were cultured in DMEM media containing 10% FBS, 100 units/ml penicillin, and 100mg/ml streptomycin and amphotericin. All cell lines were incubated at 37°C with 5% CO_2_. The cell lines were authenticated by DNA short tandem repeat (STR) profiling using Promega Geneprint 10 system in conjunction with GeneMarker HID software tool. All cell lines were made mycoplasma free if necessary with EZKill Mycoplasma removal reagent (HiMedia).

### Soft Agar assay

All experiments were performed in triplicates as described earlier (Chandrani et al, 2015). Briefly, anchorage-independent growth was assessed for the knockdown clones of *ERBB2* and *EGFR* along with respective scrambled control. About 1ml of 2X DMEM supplemented with 20% FBS containing (1ml of 1.6% agar) to obtain 0.8% agar was added to the six well plate as bottom agar and was allowed to solidify. Next, 5 * 103 cells were supplemented with 1ml of 2X DMEM containing 0.8% agar to obtain 0.4% agar and were added to the bottom agar as top agar. The cells were incubated for 2 weeks at 37°C and 5% CO2. Colonies were counted under a microscope with a magnification of 10X.

### Virus production

293FT cells were seeded in 6 well plates one day before transfection and each of the lentiviral constructs along with packaging plasmids-pPAX helper vector and pVSVG were transfected using Lipofectamine 3000 reagent (Invitrogen) as per manufacturer’s protocol. The viral soup was collected 48 and 72 hrs post transfection, passed through the 0.45µM filter and stored at 4°C. Respective cells for transduction were seeded one day before infection in a six-well plate and allowed to grow to reach 50-60% confluency. One ml of the virus soup (1:1 dilution) and 8ug/ml of polybrene (Sigma) was added to cells and incubated for six hours. Cells were selected with puromycin (Sigma) (2µg/ml) selection for 2 days as further described earlier (Upadhyay et al, 2016b).

### Growth Curve

Growth curve assay was performed on a 24 well plate format with a cell density of 20000cells/well. Cell growth was assessed post 48hr and 96hr by counting the cells using a hemocytometer and was recorded. Cell proliferation was calculated as percentage proliferation normalized to scrambled control. All the experiments were performed in triplicates.

### MTT assay

1000 cells per well were seeded in 96 well plate followed by incubation with the drug for 72 hours and six replicate per concentration and subsequently incubated with MTT (0.5 mg/ml) for 4 hours and then MTT assay was performed and data was acquired at 570nm using Microplate reader. Percentage cell viability was calculated against vehicle treated.

### Western blotting

Cells were lysed in RIPA buffer and protein concentration was estimated using BCA (MP Biomedical) method. 50 μg protein was separated on 10% SDS-PAGE gel, the transfer was verified using Ponceau S (Sigma), transferred to nitrocellulose membrane and blocked in Tris-buffered saline containing 5% BSA (Sigma) and 0.05% Tween-20(Sigma). The primary antibody against Total HER2 (sc-33684 Dilution 1:500), Total EGFR (1005) (sc-03 Dilution 1:500), Total ERK2(C-14) (sc-154 Dilution 1:500) and β-Actin(I-19)-R (sc-1616-R Dilution 1:3000) were obtained from Santa Cruz biotechnology. The primary antibodies Phospho-HER2 (Tyr1248) (AP0152 Dilution 1:500) from Abclonal and Phospho-p44/42 (T202/Y204) MAPK (#4370) Dilution 1:1000), Phospho-*EGFR* (Y1068) (#2234 Dilution 1:500) were obtained from Cell signaling technology respectively. Thiazolyl blue tetrazolium bromide (MTT, TC191) was obtained from Hi-Media.

### Receptor tyrosine kinase proteome array

The relative amount of 49 tyrosine kinases were evaluated using Proteome Profiler Human Phospho-RTK array kit (ARY001B – Proteome Profiler, R&D systems) and the protocols were followed as per manufacturer’s recommendation. Briefly, cells were harvested, washed with 1X PBS and lysed after which 400µg of protein was mixed with a buffer and incubated with pre-blocked nitrocellulose membrane at 4°C. Subsequently, the membranes were probed using detection antibodies and probed using streptavidin-HRP, after which signals were developed using the chemi-reagents provided with the kit. The Pixel density of each spot in the array in duplicate was quantified using Image J macro-Protein array analyzer plug in. The average pixel density of the duplicate spots for each of the kinases was subtracted from the negative density and was plotted, as detailed earlier (Godbole et al, 2017).

### Invasion assay

Invasion ability of the cells was assessed in Transwell system using cell culture inserts for 24 well plates with 8µm pores (BD Biosciences, NJ). The upper side of the cell culture insert was coated with Matrigel (BD Biosciences, San Jose, CA). GBC cells were seeded at a density of 2 * 10^4^ on the upper side of the coated Matrigel in presence of serum free DMEM. Complete DMEM media with 10% FBS was added to the lower side of the insert and were incubated at 37°C in 5% CO_2_ incubator for 12-14hrs. Post incubation the non-migratory cells on the lower side of the cell culture insert were removed using a cotton swab. The transwell chambers were fixed and stained with 0.1% crystal violet. The invasion ability was estimated by counting the cells that have migrated to the lower side of the cell culture insert. Cells in visual field with a magnification of 20X were counted in each Transwell chamber in triplicates.

### Wound healing assay

Confluent monolayers in 6 well plate are subjected to scratch with a sterile pipette tip. After this, cells are washed with 1X PBS to remove debris and subsequently incubated with media. Cell migration at the wound surface was measured during a period of 20h under an inverted microscope. The quantification of cell migration was done using Cell Profiler(Carpenter et al, 2006) wound healing pipeline for three independent wounds in 3 independent experiments.

### Immunohistochemistry

Immuno-histochemical analysis was done using the standard protocol of Vectastatin Universal kit. Briefly, antigen retrieval was performed by incubating the slides in pre-heated citrate buffer (pH 6) using a pressure cooker for 10 minutes. The slides were allowed to cool at room temperature before rinsing with TBST (Tris-Buffered saline-Tween 20(1%). The endogenous peroxidase activity was blocked by incubating the slides in 3% hydrogen peroxide. The slides were blocked by horse serum for 1hour before incubating with the primary antibody (HER2 DAKO A0485, Phospho-p44/42 (T202/Y204) MAPK #4370, Total ERK2(C-14) sc-154) for overnight at 4°C in moist chamber. Post incubation the slides were rinsed with TBST and incubated with universal secondary antibody (Vectastatin). The chromogenic reaction was performed using 3’-3’-diaminobenzidine chromogen solution for 5 minutes which results in brown signal. The slides are rinsed in deionized water and counterstained with hemotoxylin. Finally, the slides are dehydrated and mounted with a mounting medium and cover slip.

### Co-immunopreciptation assay

For immunoprecipitation, cells were harvested in NP-40 lysis buffer (50mM Tris pH7.4, 150mM NaCl, 0.5% NP-40, 1mM EDTA along with protease and phosphatase inhibitors), Protein lysate supernatant were combined with the anti-EGFR antibody and incubated overnight on a rotator at 4°C. Protein G-Sepharose beads (50µl) were added to the cell lysates the next day and were on a rotator at 4°C for 4 hours. The Protein G-Sepharose beads were isolated by centrifugation at 2000g for 2 minutes. Further, these beads were washed three times with NP-40 lysis buffer and heated for 10 minutes at 100 °C in loading buffer. Samples were run on SDS-PAGE and then probed by immunoblot for HER2.

### In vivo study

Five to six week old female NOD-SCID mice were injected subcutaneously with 3*106 cells/ml in 100-200 µl PBS G415 (N=13), NOZ (N=10) and OCUG1(N=10). After injecting the cells, the size of the resulting tumors was determined every third day using calipers. Afatinib inhibitor was administered to the randomized group of mice by oral gavage at 15mg/kg body weight along with vehicle control (1% Tween 80) for a period of 15 days after the tumor volume has reached between 100-150 mm3. micro PET-CT scan was performed at the end of drug treatment. The tumor volume was calculated using the formula – (Width2 * Length) /2. After 15 days, the mice were euthanized with CO2. Tumor were excised and tissues were stored for molecular and histopathological analysis.

### Statistical analysis

Prism software (GraphPad) was used to analyze proliferation and drug sensitivity of cells to inhibitors, and to determine the statistical significance of differences between the groups by applying an unpaired Student’s t test. P values < 0.05 were considered significant. The Kaplan-Meier estimation of patient survival and correlation analysis were assessed using R packages survival (http://cran.r-project.org/package=survival), and IBM SPSS v20.

## Acknowledgements

All members of the Dutt laboratory for critically reviewing the manuscript. Priyanka Bagayatkar and Vaishakhi Trivedi for *KRAS* sequencing. Medgenome Labs Pvt. Ltd for whole exome sequencing services. We thank Dr. Harsha Gowda and Dr. Akhilesh Pandey, IOB Bangalore for sharing the gallbladder cancer cell lines; the BTIS facility, funded by the Department of Biotechnology (DBT), Govt. of India, at ACTREC; and, the in-house Genomics facility for Sanger Sequencing at ACTREC. A.D. is supported by an Intermediate Fellowship from the Wellcome Trust/DBT India Alliance (IA/I/11/2500278). P.I. is supported by a Senior Research Fellowship from ACTREC. The funders had no role in study design, data collection and analysis, decision to publish or preparation of the manuscript

## Author contributions

P.I., and A.D. designed research. P.I., M.R., P.C., N.G., B.D., S.S., R.P., R.T., B. M., and B.S. performed research. S.G.B., V.C., A.C., M.R.R., K.P., S.D. and S.V.S. contributed reagents and samples. P.I., M.R., P.C. N.G., A.J., H.K., P. Chau., A.I., and A.D. analyzed data. P.I. and A.D. wrote the paper.

## The Paper Explained

### Problem

Are there somatic alterations that might serve as promising therapeutic targets in early-stage gallbladder tumors?

### Results

Findings from this study implicate *ERBB2* as an important therapeutic target in early stage gallbladder cancer. We also present the first evidence that the presence of *KRAS* (G12V), but not *KRAS* (G13D) mutation, may preclude gallbladder cancer patients to respond to anti-EGFR treatment.

### Impact

This study could lead to the adoption of an approved clinical algorithm, commonly practiced to opt for anti-EGFR treatment in colorectal cancer, to treat gallbladder cancer patients.

## References and Citations

1. Barreto SG, Dutt A, Chaudhary A (2014) A genetic model for gallbladder carcinogenesis and its dissemination. Annals of oncology: official journal of the European Society for Medical Oncology 25: 1086–1097

2. Boeva V, Popova T, Bleakley K, Chiche P, Cappo J, Schleiermacher G, Janoueix-Lerosey I, Delattre O, Barillot E (2012) Control-FREEC: a tool for assessing copy number and allelic content using next-generation sequencing data. Bioinformatics 28: 423–425

3. Bose R, Kavuri SM, Searleman AC, Shen W, Shen D, Koboldt DC, Monsey J, Goel N, Aronson AB, Li S, Ma CX, Ding L, Mardis ER, Ellis MJ (2013) Activating HER2 mutations in HER2 gene amplification negative breast cancer. Cancer discovery 3: 224–237

4. Carpenter AE, Jones TR, Lamprecht MR, Clarke C, Kang IH, Friman O, Guertin DA, Chang JH, Lindquist RA, Moffat J, Golland P, Sabatini DM (2006) CellProfiler: image analysis software for identifying and quantifying cell phenotypes. Genome biology 7: R100

5. Chandrani P, Prabhash K, Prasad R, Sethunath V, Ranjan M, Iyer P, Aich J, Dhamne H, Iyer DN, Upadhyay P, Mohanty B, Chandna P, Kumar R, Joshi A, Noronha V, Patil V, Ramaswamy A, Karpe A, Thorat R, Chaudhari P, Ingle A, Choughule A, Dutt A (2017) Drug-sensitive FGFR3 mutations in lung adenocarcinoma. Annals of oncology: official journal of the European Society for Medical Oncology 28: 597–603

6. Chandrani P, Upadhyay P, Iyer P, Tanna M, Shetty M, Raghuram GV, Oak N, Singh A, Chaubal R, Ramteke M, Gupta S, Dutt A (2015) Integrated genomics approach to identify biologically relevant alterations in fewer samples. BMC genomics 16: 936

7. Choughule A, Sharma R, Trivedi V, Thavamani A, Noronha V, Joshi A, Desai S, Chandrani P, Sundaram P, Utture S, Jambhekar N, Gupta S, Aich J, Prabhash K, Dutt A (2014) Coexistence of KRAS mutation with mutant but not wild-type EGFR predicts response to tyrosine-kinase inhibitors in human lung cancer. British journal of cancer 111: 2203–2204

8. Christensen JG, Zou HY, Arango ME, Li Q, Lee JH, McDonnell SR, Yamazaki S, Alton GR, Mroczkowski B, Los G (2007) Cytoreductive antitumor activity of PF-2341066, a novel inhibitor of anaplastic lymphoma kinase and c-Met, in experimental models of anaplastic large-cell lymphoma. Molecular cancer therapeutics 6: 3314–3322

9. De Roock W, Claes B, Bernasconi D, De Schutter J, Biesmans B, Fountzilas G, Kalogeras KT, Kotoula V, Papamichael D, Laurent-Puig P, Penault-Llorca F, Rougier P, Vincenzi B, Santini D, Tonini G, Cappuzzo F, Frattini M, Molinari F, Saletti P, De Dosso S, Martini M, Bardelli A, Siena S, Sartore-Bianchi A, Tabernero J, Macarulla T, Di Fiore F, Gangloff AO, Ciardiello F, Pfeiffer P, Qvortrup C, Hansen TP, Van Cutsem E, Piessevaux H, Lambrechts D, Delorenzi M, Tejpar S (2010a) Effects of KRAS, BRAF, NRAS, and PIK3CA mutations on the efficacy of cetuximab plus chemotherapy in chemotherapy-refractory metastatic colorectal cancer: a retrospective consortium analysis. The Lancet Oncology 11: 753–762

10. De Roock W, Jonker DJ, Di Nicolantonio F, Sartore-Bianchi A, Tu D, Siena S, Lamba S, Arena S, Frattini M, Piessevaux H, Van Cutsem E, O’Callaghan CJ, Khambata-Ford S, Zalcberg JR, Simes J, Karapetis CS, Bardelli A, Tejpar S (2010b) Association of KRAS p.G13D mutation with outcome in patients with chemotherapy-refractory metastatic colorectal cancer treated with cetuximab. Jama 304: 1812–1820

11. Do K, O’Sullivan Coyne G, Chen AP (2015) An overview of the NCI precision medicine trials-NCI MATCH and MPACT. Chinese clinical oncology 4: 31

12. Doval DC, Azam S, Sinha R, Batra U, Mehta A (2014) Expression of epidermal growth factor receptor, p53, Bcl2, vascular endothelial growth factor, cyclooxygenase-2, cyclin D1, human epidermal receptor-2 and Ki-67: Association with clinicopathological profiles and outcomes in gallbladder carcinoma. J Carcinog 13: 10

13. Druker BJ, Sawyers CL, Kantarjian H, Resta DJ, Reese SF, Ford JM, Capdeville R, Talpaz M (2001) Activity of a specific inhibitor of the BCR-ABL tyrosine kinase in the blast crisis of chronic myeloid leukemia and acute lymphoblastic leukemia with the Philadelphia chromosome. The New England journal of medicine 344: 1038–1042

14. Flaherty KT, Puzanov I, Kim KB, Ribas A, McArthur GA, Sosman JA, O’Dwyer PJ, Lee RJ, Grippo JF, Nolop K, Chapman PB (2010) Inhibition of mutated, activated BRAF in metastatic melanoma. The New England journal of medicine 363: 809–819

15. Forbes SA, Bhamra G, Bamford S, Dawson E, Kok C, Clements J, Menzies A, Teague JW, Futreal PA, Stratton MR (2008) The Catalogue of Somatic Mutations in Cancer (COSMIC). Current protocols in human genetics Chapter 10: Unit 10 11

16. Garner AP, Bialucha CU, Sprague ER, Garrett JT, Sheng Q, Li S, Sineshchekova O, Saxena P, Sutton CR, Chen D, Chen Y, Wang H, Liang J, Das R, Mosher R, Gu J, Huang A, Haubst N, Zehetmeier C, Haberl M, Elis W, Kunz C, Heidt AB, Herlihy K, Murtie J, Schuller A, Arteaga CL, Sellers WR, Ettenberg SA (2013) An antibody that locks HER3 in the inactive conformation inhibits tumor growth driven by HER2 or neuregulin. Cancer research 73: 6024–6035

17. Godbole M, Tiwary K, Badwe R, Gupta S, Dutt A (2017) Progesterone suppresses the invasion and migration of breast cancer cells irrespective of their progesterone receptor status - a short report. Cellular oncology 40: 411–417

18. Gonzalez-Perez A, Perez-Llamas C, Deu-Pons J, Tamborero D, Schroeder MP, Jene-Sanz A, Santos A, Lopez-Bigas N (2013) IntOGen-mutations identifies cancer drivers across tumor types. Nature methods 10: 1081–1082

19. Iyer P, Barreto SG, Sahoo B, Chandrani P, Ramadwar MR, Shrikhande SV, Dutt A (2016) Non-typhoidal Salmonella DNA traces in gallbladder cancer. Infectious agents and cancer 11: 12

20. Javle M, Churi C, Kang HC, Shroff R, Janku F, Surapaneni R, Zuo M, Barrera C, Alshamsi H, Krishnan S, Mishra L, Wolff RA, Kaseb AO, Thomas MB, Siegel AB (2015) HER2/neu-directed therapy for biliary tract cancer. Journal of hematology & oncology 8: 58

21. Jiao Y, Pawlik TM, Anders RA, Selaru FM, Streppel MM, Lucas DJ, Niknafs N, Guthrie VB, Maitra A, Argani P, Offerhaus GJA, Roa JC, Roberts LR, Gores GJ, Popescu I, Alexandrescu ST, Dima S, Fassan M, Simbolo M, Mafficini A, Capelli P, Lawlor RT, Ruzzenente A, Guglielmi A, Tortora G, de Braud F, Scarpa A, Jarnagin W, Klimstra D, Karchin R, Velculescu VE, Hruban RH, Vogelstein B, Kinzler KW, Papadopoulos N, Wood LD (2013) Exome sequencing identifies frequent inactivating mutations in BAP1, ARID1A and PBRM1 in intrahepatic cholangiocarcinomas. Nature genetics 45: 1470–1473

22. Kandioler-Eckersberger D, Ludwig C, Rudas M, Kappel S, Janschek E, Wenzel C, Schlagbauer-Wadl H, Mittlbock M, Gnant M, Steger G, Jakesz R (2000) TP53 mutation and p53 overexpression for prediction of response to neoadjuvant treatment in breast cancer patients. Clinical cancer research: an official journal of the American Association for Cancer Research 6: 50–56

23. Krause DS, Van Etten RA (2005) Tyrosine kinases as targets for cancer therapy. The New England journal of medicine 353: 172–187

24. Kumar SS, Price TJ, Mohyieldin O, Borg M, Townsend A, Hardingham JE (2014) KRAS G13D Mutation and Sensitivity to Cetuximab or Panitumumab in a Colorectal Cancer Cell Line Model. Gastrointestinal cancer research: GCR 7: 23–26

25. Lawrence MS, Stojanov P, Polak P, Kryukov GV, Cibulskis K, Sivachenko A, Carter SL, Stewart C, Mermel CH, Roberts SA, Kiezun A, Hammerman PS, McKenna A, Drier Y, Zou L, Ramos AH, Pugh TJ, Stransky N, Helman E, Kim J, Sougnez C, Ambrogio L, Nickerson E, Shefler E, Cortes ML, Auclair D, Saksena G, Voet D, Noble M, DiCara D, Lin P, Lichtenstein L, Heiman DI, Fennell T, Imielinski M, Hernandez B, Hodis E, Baca S, Dulak AM, Lohr J, Landau DA, Wu CJ, Melendez-Zajgla J, Hidalgo-Miranda A, Koren A, McCarroll SA, Mora J, Lee RS, Crompton B, Onofrio R, Parkin M, Winckler W, Ardlie K, Gabriel SB, Roberts CW, Biegel JA, Stegmaier K, Bass AJ, Garraway LA, Meyerson M, Golub TR, Gordenin DA, Sunyaev S, Lander ES, Getz G (2013) Mutational heterogeneity in cancer and the search for new cancer-associated genes. Nature 499: 214–218

26. Le Tourneau C, Paoletti X, Servant N, Bieche I, Gentien D, Rio Frio T, Vincent-Salomon A, Servois V, Romejon J, Mariani O, Bernard V, Huppe P, Pierron G, Mulot F, Callens C, Wong J, Mauborgne C, Rouleau E, Reyes C, Henry E, Leroy Q, Gestraud P, La Rosa P, Escalup L, Mitry E, Tredan O, Delord JP, Campone M, Goncalves A, Isambert N, Gavoille C, Kamal M (2014) Randomised proof-of-concept phase II trial comparing targeted therapy based on tumour molecular profiling vs conventional therapy in patients with refractory cancer: results of the feasibility part of the SHIVA trial. British journal of cancer 111: 17–24

27. Li D, Ambrogio L, Shimamura T, Kubo S, Takahashi M, Chirieac LR, Padera RF, Shapiro GI, Baum A, Himmelsbach F, Rettig WJ, Meyerson M, Solca F, Greulich H, Wong KK (2008) BIBW2992, an irreversible EGFR/HER2 inhibitor highly effective in preclinical lung cancer models. Oncogene 27: 4702–4711

28. Li M, Zhang Z, Li X, Ye J, Wu X, Tan Z, Liu C, Shen B, Wang XA, Wu W, Zhou D, Zhang D, Wang T, Liu B, Qu K, Ding Q, Weng H, Ding Q, Mu J, Shu Y, Bao R, Cao Y, Chen P, Liu T, Jiang L, Hu Y, Dong P, Gu J, Lu W, Shi W, Lu J, Gong W, Tang Z, Zhang Y, Wang X, Chin YE, Weng X, Zhang H, Tang W, Zheng Y, He L, Wang H, Liu Y, Liu Y (2014) Whole-exome and targeted gene sequencing of gallbladder carcinoma identifies recurrent mutations in the ErbB pathway. Nature genetics 46: 872–876

29. Li W, Qiu T, Zhi W, Shi S, Zou S, Ling Y, Shan L, Ying J, Lu N (2015) Colorectal carcinomas with KRAS codon 12 mutation are associated with more advanced tumor stages. BMC cancer 15: 340

30. Linggi B, Carpenter G (2006) ErbB receptors: new insights on mechanisms and biology. Trends Cell Biol 16:649–656

31. McGranahan N, Swanton C (2015) Biological and therapeutic impact of intratumor heterogeneity in cancer evolution. Cancer Cell 27: 15–26

32. Miller G, Jarnagin WR (2008) Gallbladder carcinoma. Eur J Surg Oncol 34: 306–312

33. Muller BG, De Aretxabala X, Gonzalez Domingo M (2014) A review of recent data in the treatment of gallbladder cancer: what we know, what we do, and what should be done. American Society of Clinical Oncology educational book American Society of Clinical Oncology Meeting: e165–170

34. Nakamura H, Arai Y, Totoki Y, Shirota T, Elzawahry A, Kato M, Hama N, Hosoda F, Urushidate T, Ohashi S, Hiraoka N, Ojima H, Shimada K, Okusaka T, Kosuge T, Miyagawa S, Shibata T (2015) Genomic spectra of biliary tract cancer. Nature genetics 47: 1003–1010

35. Osumi H, Shinozaki E, Osako M, Kawazoe Y, Oba M, Misaka T, Goto T, Kamo H, Suenaga M, Kumekawa Y, Ogura M, Ozaka M, Matsusaka S, Chin K, Hatake K, Mizunuma N (2015) Cetuximab treatment for metastatic colorectal cancer with KRAS p.G13D mutations improves progression-free survival. Molecular and clinical oncology 3: 1053–1057

36. Paez JG, Janne PA, Lee JC, Tracy S, Greulich H, Gabriel S, Herman P, Kaye FJ, Lindeman N, Boggon TJ, Naoki K, Sasaki H, Fujii Y, Eck MJ, Sellers WR, Johnson BE, Meyerson M (2004) EGFR mutations in lung cancer: correlation with clinical response to gefitinib therapy. Science 304: 1497–1500

37. Petitjean A, Achatz MI, Borresen-Dale AL, Hainaut P, Olivier M (2007) TP53 mutations in human cancers: functional selection and impact on cancer prognosis and outcomes. Oncogene 26: 2157–2165

38. Prior IA, Lewis PD, Mattos C (2012) A comprehensive survey of Ras mutations in cancer. Cancer research 72: 2457–2467

39. Rakic M, Patrlj L, Kopljar M, Klicek R, Kolovrat M, Loncar B, Busic Z (2014) Gallbladder cancer. Hepatobiliary surgery and nutrition 3: 221–226

40. Ramos AH, Lichtenstein L, Gupta M, Lawrence MS, Pugh TJ, Saksena G, Meyerson M, Getz G (2015) Oncotator: cancer variant annotation tool. Human mutation 36: E2423–2429

41. Rusnak DW, Lackey K, Affleck K, Wood ER, Alligood KJ, Rhodes N, Keith BR, Murray DM, Knight WB, Mullin RJ, Gilmer TM (2001) The effects of the novel, reversible epidermal growth factor receptor/ErbB-2 tyrosine kinase inhibitor, GW2016, on the growth of human normal and tumor-derived cell lines in vitro and in vivo. Molecular cancer therapeutics 1: 85–94

42. Sherry ST, Ward MH, Kholodov M, Baker J, Phan L, Smigielski EM, Sirotkin K (2001) dbSNP: the NCBI database of genetic variation. Nucleic acids research 29: 308–311

43. Spivak-Kroizman T, Rotin D, Pinchasi D, Ullrich A, Schlessinger J, Lax I (1992) Heterodimerization of c-erbB2 with different epidermal growth factor receptor mutants elicits stimulatory or inhibitory responses. The Journal of biological chemistry 267: 8056–8063

44. Tebbutt N, Pedersen MW, Johns TG (2013) Targeting the ERBB family in cancer: couples therapy. Nature reviews Cancer 13: 663–673

45. Upadhyay P, Gardi N, Desai S, Sahoo B, Singh A, Togar T, Iyer P, Prasad R, Chandrani P, Gupta S, Dutt A (2016a) TMC-SNPdb: an Indian germline variant database derived from whole exome sequences. Database: the journal of biological databases and curation 2016

46. Upadhyay P, Nair S, Kaur E, Aich J, Dani P, Sethunath V, Gardi N, Chandrani P, Godbole M, Sonawane K, Prasad R, Kannan S, Agarwal B, Kane S, Gupta S, Dutt S, Dutt A (2016b) Notch pathway activation is essential for maintenance of stem-like cells in early tongue cancer. Oncotarget 7: 50437–50449

47. Yanagisawa N, Yamashita K, Kuba T, Okayasu I (2010) Sporadic TP53 transition mutations in chronic cholecystitis are possibly linked to gallbladder carcinogenesis. Anticancer research 30: 4443–4449

48. Yoon HH, Tougeron D, Shi Q, Alberts SR, Mahoney MR, Nelson GD, Nair SG, Thibodeau SN, Goldberg RM, Sargent DJ, Sinicrope FA, Alliance for Clinical Trials in O (2014) KRAS codon 12 and 13 mutations in relation to disease-free survival in BRAF-wild-type stage III colon cancers from an adjuvant chemotherapy trial (N0147 alliance). Clinical cancer research: an official journal of the American Association for Cancer Research 20: 3033–3043

49. Yoshida K, Ogawa S (2014) Splicing factor mutations and cancer. Wiley interdisciplinary reviews RNA 5: 445–459

50. Zhou X, Agazie YM (2012) The signaling and transformation potency of the overexpressed HER2 protein is dependent on the normally-expressed EGFR. Cellular signalling 24: 140–150

51. Zhu AX, Hong TS, Hezel AF, Kooby DA (2010) Current management of gallbladder carcinoma. Oncologist 15: 168–181

